# The mycotoxin Beauvericin is an uncompetitive inhibitor of Cathepsin B

**DOI:** 10.1101/2025.01.17.633572

**Authors:** Xiaoli Yang, Pablo Cea-Medina, Mohanraj Gopalswamy, Aparna Vaidya, Sonja Schavier, Shixin Oltzen, Sofie Moßner, Anfei Huang, Jing Qi, Doreen Floss, Markus Uhrberg, Holger Gohlke, Stefanie Scheu

## Abstract

Beauvericin (BEA), a cyclic depsipeptide, is a mycotoxin of the enniatin family and the secondary metabolite of various toxigenic fungi. Multiple biological functions of BEA have been well investigated, such as anti-cancer, anti-inflammatory, anti-microbial, and immune-activating functions. In a recent study, we showed that BEA can target Toll-like receptor 4 (TLR4) to induce dendritic cell (DC) activation. In an in silico screen, we identified Cathepsin B (CTSB) as a potential additional interaction partner for BEA, which has been verified recently in a study showing inhibition of CTSB activity by BEA in cell-free assays. The underlying molecular mechanism of BEA-mediated CTSB inhibition remains unknown as do the cellular entities where this inhibition takes place. In this study, we determine the effects of BEA on CTSB within granulocyte-macrophage colony-stimulating factor (GM-CSF)-cultured bone marrow-derived dendritic cells (BMDCs) and human leukemia monocytic cell line THP-1 induced dendritic cells (iDCs). BEA significantly suppresses CTSB activity in both mouse BMDCs and human iDCs. NMR analyses indicate that BEA directly interacts with CTSB. Enzyme kinetics show that BEA can directly inhibit CTSB activity and acts as an uncompetitive inhibitor. Molecular docking analysis revealed a putative binding site for BEA in human CTSB. Collectively, we describe the molecular mechanisms underlying the biological activity of BEA against human CTSB, suggesting that CTSB may be a candidate target for tumor therapy.

## Introduction

CTSB is a member of the lysosomal cysteine protease family, which also includes cathepsin C, F, H, K, L, O, S, V, X, and W (1, 2). Structurally, CTSB has the regular V-shaped active site cleft of papain-like enzymes formed by the folding of two distinct domains. In addition, CTSB has a 20-residue insertion, termed occluding loop, that overlaps with the active site, binds the C-terminus of the protein, and induces carboxypeptidase activity (1, 3). Unlike other members of the papain family, CTSB has both exopeptidase and endopeptidase activity. Its exopeptidase activity is dependent on the stability of the occluding loop, which is controlled by acidic pH. By contrast, the endopeptidase activity of CTSB is determined by neutral/alkaline pH, which can disrupt the salt bridges between the occluding loop and lead to the exposure of the active site (3, 4).

CTSB has been shown to play an important role in intracellular proteolysis and remodeling of the extracellular matrix of cellular proteolysis networks (3, 4). Under normal physiological conditions, CTSB activity is well controlled and has been implicated in several oncogenic processes such as bone resorption, antigen processing, and protein turnover (5). However, aberrant expression of the cysteine protease CTSB is observed in the pathogenesis of many diseases, including neurological (6–8), immune (9–11), and cardiac diseases (12, 13) as well as cancer (14, 15). Numerous studies have shown that CTSB overexpression is correlated with invasive and metastatic cancers because it supports a pro-malignant phenotype by enabling the maintenance of active proteases through active suppression of their inhibitors (16–18). For example, CTSB enhances the activity of matrix metalloproteinases (MMPs) by destroying their inhibitors (e.g., TIMP1 and TIMP2) in human articular chondrocytes and maintaining high levels of MMPs, thereby promoting extracellular matrix (ECM) degradation and angiogenesis (19). Therefore, targeting CTSB could have significant clinical relevance in the treatment of various cancers.

BEA is a cyclic hexadepsipeptide (20), composed of three units of phenylalanine (Phe) and three units of 2-hydroxyisovaleric acid (Hiv), and is produced by various fungi, such as *Beaveria bassiana* and *Fusarium spp* (21, 22). The bioactivity of BEA has been studied in the context of anti-cancer (23, 24), anti-virus (25, 26), anti-inflammatory (27, 28), antimicrobial (29–32), immune-regulating (33), insecticidal (34, 35), and pesticidal activity (36, 37). We as well as others have identified CTSB as a molecular target of BEA (38). However, the mode of action of BEA binding to CTSB as well as if this also takes place in treatment-relevant immune cells remains unclear. In this study, BEA is shown to be an uncompetitive inhibitor of CTSB in mouse and human DCs. In addition, the binding site of BEA in CTSB was revealed using a molecular docking approach.

## Materials and Methods

### Mice

Cells from the bone marrow of C57BL/6N mice were used for GM-CSF culture. No experiments on live animals were performed. Mice were euthanized by cervical dislocation before bone marrow was harvested. The euthanasia method used is in strict accordance with accepted norms of veterinary best practice. Animals were kept under specific pathogen-free conditions in the animal research facility of the Heinrich Heine University of Düsseldorf strictly according to German animal welfare guidelines.

### BMDC culture and stimulation conditions

2x10^6^ mouse bone marrow cells were cultured in non-treated 94x16 mm dishes (Sarstedt, Cat# 82.1473) in 10 ml VLE DMEM (Biochrom, Cat# P04-04515) containing 10% heat-inactivated FCS (Sigma-Aldrich, Cat# F7524), 0.1% 2-mercaptoethanol (Thermo Fischer Scientific, Cat# 31350-010), and GM-CSF and kept for 9 days. GM-CSF cultures were performed as previously described (39). 10 ml GM-CSF containing medium was added to the plates on day 3. On day 6, 10 ml medium was carefully removed and centrifuged. The cell pellet was resuspended in 10 ml medium and added to the dish. On day 9, BMDCs were seeded on a 24-well plate and subjected to a specified BEA concentration or to 30 μM of the control inhibitor, CA074 (MedChemExpress, Cat# HY-103350), for 24 hours. The stimulated cells were collected for FACS or CTSB activity assays.

### iDC culture and stimulation conditions

iDCs were generated according to the protocol (40), namely, 1x10^6^ THP1 cells were cultured in a T25 flask with 5 ml RPMI (Thermo Fisher Scientific, Cat# 31870-025) including 10% heat-inactivated FCS (Sigma-Aldrich, Cat# F7524), 0.1% 2-mercaptoethanol (Thermo Fischer Scientific, Cat# 31350-010), 1% PenStrep (Biochrom GmbH, Cat#A2212), 1500 IU/ml rhGM-CSF (ImmunoTools, Cat#11343125), and 1500 IU/ml rhIL-4 (ImmunoTools, Cat#11340045) and incubated at 37°C and 5% CO_2_ for 5 days. On day 3, 4 ml of the medium was exchanged for fresh medium including 1500 IU/ml rhGM-CSF (ImmunoTools, Cat#11343125) and 1500 IU/ml rhIL-4 (ImmunoTools, Cat#11340045). On day 5, cells are seeded on a 24-well plate and subjected to the specified concentration of BEA and CA074 (30 μM) (MedChemExpress, Cat# HY-103350) for 24 hours; the stimulated cells were collected for FACS or CTSB activity assays.

### Cell-based CTSB activity assays

The stimulated cells were lysed with chilled cell lysis buffer (Abcam, Cat# ab65300), and the cell lysates were subjected to protein quantification by BCA. Cell CTSB activity was measured according to the manufacturer’s protocol (Abcam, Cat# ab65300). Briefly, equal amounts of protein in 50 µl reaction buffer were added to 96 black clear bottom-well plates (Greiner Bio-One, Cat# 675086), followed by the addition of 50 µl cathepsin B reaction buffer. The reaction was then incubated at 37°C for 1 hour with the addition of substrate and protected from light. The output was read on a fluorescence microplate reader at Ex/Em = 400/505 nm.

Alternatively, stimulated cells were incubated with Rhodamine 110-(RR)2 substrate (Abcam, Cat# ab270787) at approximately 1/50 for 30 minutes at 37°C, protected from light. After incubation, the cells were stained with Hoechst 33342 (Abcam, Cat# ab270787) at approximately 0.5% for 10 minutes and then analyzed by FACS.

### STD-NMR analysis

NMR experiments were performed at 298 K on Bruker Avance III HD spectrometers operating at 750 MHz, equipped with 5 mm triple resonance TCI (^1^H, ^13^C, ^15^N) cryoprobes and shielded z-gradients. BEA compound was purchased (Item No. 11426) from Cayman Chemical, Ann Arbor, Michigan, USA, and recombinant human CTSB (6His tag at the C-terminus) was purchased (Item No. 153921.10) from Biomol GmbH, Hamburg, Germany. Both CTSB and BEA were used without any further purifications. The purity of the compound (>95%) was checked by 1D ^1^H NMR of 250 µM BEA compound measured (256 scans) in 90% (v/v) DMSO_d6_. (Figure 2A). BEA is sparingly soluble in aqueous buffers, so 0.1% (v/v) Tween 20 was used to increase the compound solubility. To perform STD-NMR experiments, 5 µM of CTSB and 250 µM BEA were prepared in 25 mM oxalic acid/sodium oxalate, 5% (v/v) DMSO_d6_, pH 5.0, 10% (v/v) D_2_O, and 0.1% (v/v) Tween 20. The STD-NMR spectra were acquired with 4928 scans, and the subtraction of the reference spectrum was performed internally via phase cycling after every scan to obtain the STD effect. Selective saturation of protein resonances (on-resonance spectrum) was performed by irradiating at -0.3 ppm for a total saturation time of 3 s with a delay time (D1) of 5 s. There was no compound signal detected below 0 ppm. For the reference spectrum (off-resonance), the samples were irradiated at −40 ppm. The STD intensity of the aromatic proton STD effect was set to 100% as a reference, and the relative intensities were determined (41). The assignment of the protons of the ligand was achieved by the analysis of the 1D ^1^H NMR and aided by ^1^H chemical shift prediction performed by the ChemNMR package, contained within the software ChemDraw (PerkinElmer). Data was processed and analyzed with TopSpin 3.2 (Bruker BioSpin). Sodium 2,2-dimethyl-2-silapentane-5-sulfonate (DSS) was used for chemical shift referencing.

### Enzyme kinetics

Recombinant human CTSB (R&D Biosystems, Cat# 953-CY) was activated in a solution of 25 mM MES pH 5 and 5 mM DTT at a protein concentration of 25 µg/ml, as indicated by the manufacturer. The activation was carried out by incubating the solution in a thermal bath at 26 °C for 30 minutes and then diluting to a concentration of 0.2 µg/ml in a 25 mM MES buffer pH 5.

Reactions were carried out in a 96-Well black microplate (Greiner Bio-One, Cat# 675086) in a final volume of 100 µl, using 20 ng of enzyme with increasing concentrations of the fluorogenic substrate Z-Leu-Arg AMC (R&D Biosystems, Cat# ES008) (from 0 to 200 µM) and BEA (from 0 to 25 µM).

Progress curves were measured using a TECAN infinite M plex microplate reader with an excitation wavelength of 380 nm and an emission wavelength of 460 nm. All the components, except the enzyme, were loaded onto the plate and were left to reach a temperature of 26 °C. Protein samples were loaded using a multichannel pipette. Plates were shaken using a pulse of 3.5 s with an amplitude of 4.5 mm. Data was collected for a total of 1400 s with a reading interval of 45 s.

Reaction rates were calculated from the progress curves using linear regression. Kinetic parameters were derived by fitting a Michaelis-Menten model to a plot of reaction rates at different substrate concentrations as described in equation (1)

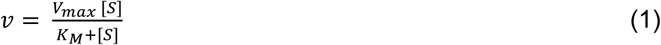

where [*S*] is the substrate concentration, *v* is the observed reaction rate or velocity, *V*_max_ is the maximal velocity of the enzyme, and *K*_M_ is the Michaelis-Menten constant. To later derive an inhibition constant, we employed a general inhibitor model (eq. (2)), as described by Cornish-Bowden in ref. (42)

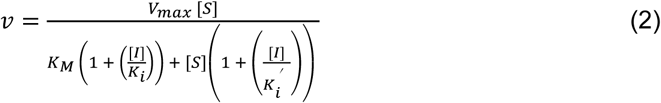

where [*I*] is the concentration of the inhibitor, *K*_i_ is the inhibition constant of the [EI] complex and *K*_i_’ is the inhibition constant for the [ESI] complex. The equation can be reformulated as shown in equation (3)

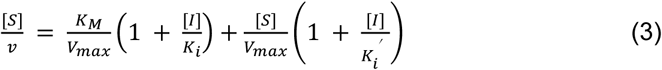

which allows us to derive the *K*_i_ of competitive, non-competitive, and uncompetitive inhibitors. The interception point of curves at different [*S*] on the x-axis is:

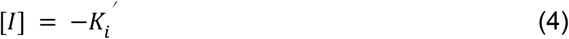

and the value on the y-axis for the intercept is:

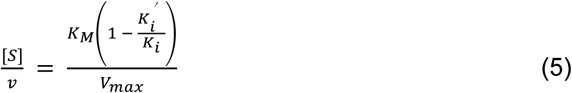

For uncompetitive inhibitors the value of *K*_i_ → ∞, therefore the y-value of the intercept is:

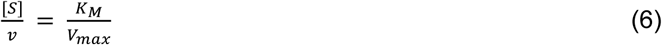

To approximate the observed interception point, we identified the (x, y)-values that minimize the sum of the squared distances perpendicular to each substrate curve *i*

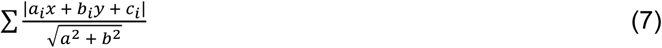

where *a*_i_, *b*_i_, and *c*_i_ correspond to the coefficients of the curves at different [*S*], derived by fitting to equation (3).

All data manipulation, including model fitting, was performed using the Pandas, NumPy, and SciPy modules of Python 3.10.

### Modeling and co-solvent probe molecular dynamics simulations of human CTSB

Given the high degree of conformational variability shown by human CTSB, we employed four crystal structures as starting points: PDB ID 1CSB, 3CBK, 1HUC, and 3K9M. To avoid biasing the results due to differences in the sequence length of each solved crystal structure, and to fix missing loops and residues, all the structures were remodeled using Modeller (43), using as a target the sequence of residues Phe74 to Ile339 of human CTSB (UniProtKB ID: P07858) and the respective crystal structure as a template. 30 models were generated for each system, and the best one was selected based on the DOPE score for further work.

Each system was packed in a solvent box with either benzene or isopropanol as co-solvent using PACKMOL-Memgen (44). A ratio of 99 water molecules per co-solvent molecule (benzene or isopropanol) was utilized. Five different random starting configurations of co-solvent distribution were generated for each system. The most likely protonation state for titratable residues was assigned with Propka (45). Parameter files were generated using tleap in AmberTools (46), with ff19SB (47) as protein force field and the co-solvent parameters as distributed alongside PACKMOL-Memgen, TIP3P as water model (48) and K^+^ and Cl^-^ counter ions with parameters from Joung & Cheatham (49). All simulations were performed using *pmemd.cuda* on Amber22 (50), using a direct space cut-off of 10 Å for long-range electrostatic interactions. All bonds involving hydrogen atoms were constrained using the SHAKE algorithm (51).

The systems were minimized in three consecutive rounds; first using positional restraint forces of 5 kcal mol^-1^ Å^-2^, followed by restraint forces of 0.1 kcal mol^-1^ Å^-2^, and lastly a restraint-free minimization. Each minimization round was carried out using 500 steps of steepest descent followed by 2000 steps of conjugate gradient. The systems were thermalized from 0 to 100 K in a window of 25 ps, using the Langevin thermostat with a collision coefficient of 1.0 ps^-1^ under NVT conditions. Afterward, the systems were heated to 300 K over a window of 250 ps under NPT conditions using the Berendsen barostat with isotropic position scaling. Both thermalization and pressure adaptation were carried out using positional restraint forces of 1.0 kcal mol^-1^ Å^-2^. Five subsequent 50 ps-long NPT runs were carried out, reducing the positional restraint forces by 0.2 kcal mol^-1^ Å^-2^ at each step until reaching zero. Two production runs of 600 ns length were performed for each system using a time step of 2 fs and saving coordinates every 100 ps. This yielded a cumulative simulation time of 1.2 µs per starting configuration. Since five random initial solvent configurations were generated for each co-solvent, the total cumulative sampling time for each co-solvent probe was 6 µs. All simulations were analyzed using *cpptraj* (52). The protein was aligned to the first frame of each trajectory with the *rmsd* command, using the backbone atoms. The density of each co-solvent was calculated using the *grid* command with a spacing of 0.5 Å.

### Docking of BEA against the inactive conformation of human CTSB

Molecular docking of BEA was performed against the inactive conformation of human CTSB (PDB ID 3PBH), with the propeptide region removed. The protein structure was prepared using the protein preparation wizard, with protonation states assigned by Propka, while the ligand was prepared with LigPrep (LigPrep, Schrödinger, LLC). A grid with dimensions of 35 Å was centered at the midpoint of residues Phe30 and Val33 of the removed propeptide.

Due to the large size of the ligand, a rigid docking using a diverse conformational ensemble of BEA structures was used. The conformational ensemble was generated using Schrödinger’s macrocycle conformational sampling tool, using an RMSD cut-off of 0.75 Å and default options (53). 255 non-redundant conformations were generated.

Rigid dockings were carried out using Glide in extra precision mode (XP) (54) using OPLS_2005 as a force field. Three poses were generated for each conformation of BEA. The resulting binding modes were analyzed using PyMOL.

### Statistical analysis

The GraphPad Prism 9.0 software was used for data analysis. Data are represented as means ± SEM. For analyzing statistical significance between multiple groups, a one-way ANOVA with Dunnett’s multiple comparisons test was used. For analyzing statistical significance for comparisons of more than two groups with two or more stimulations, two-way ANOVA with Sidak’s multiple comparisons test was used. All *p* values < 0.05 were considered statistically significant.

## Results

### BEA suppresses CTSB activity in DCs

Our previous study showed that BEA can target TLR4 to induce DC activation (33). In searching for alternative molecular targets of BEA in DCs, we performed in silico studies and found CTSB as a potential target of BEA. Recently, Silva et al. indicated that BEA potently inhibits human CTSB activity in cell-free assays (38). We wondered if BEA could inhibit CTSB in immune cells relevant for anti-cancer immunotherapy approaches such as DCs. To this end, BMDCs were generated by culturing bone marrow cells with GM-CSF followed by stimulation with BEA at various concentrations for 16 hours. First, the CTSB enzyme activity of BEA-stimulated BMDCs was determined in the cell lysates. CTSB activity was significantly suppressed by BEA stimulation already at 2.5 µM showing a dose-dependent inhibition of CTSB activity and suggesting that CTSB is a target of BEA in BMDCs (**Figure 1A**). CTSB activity was also detected in cells using a flow cytometry-based assay. Consistently, BEA can inhibit CTSB activity in BMDCs, and significant inhibition was observed starting at 7.5 µM BEA in this assay (**Figure 1B and 1C**). This led us to ask whether BEA could also inhibit human CTSB. To address this question, human monocyte-derived immature dendritic cells (iDCs) were induced from THP-1 cells in the presence of human GM-CSF and IL-4 and then subjected to various concentrations of BEA stimulation for 16 hours. As shown in **Figure 1D**, BEA can also suppress human CTSB activity with significant inhibition observed starting at a concentration 5 µM BEA. In addition, flow cytometry data also show that BEA significantly inhibits human CTSB activity (**Figure 1E and 1F**). Taken together, BEA inhibits CTSB activity in mouse BMDCs and human iDCs.

**Fig 1.**
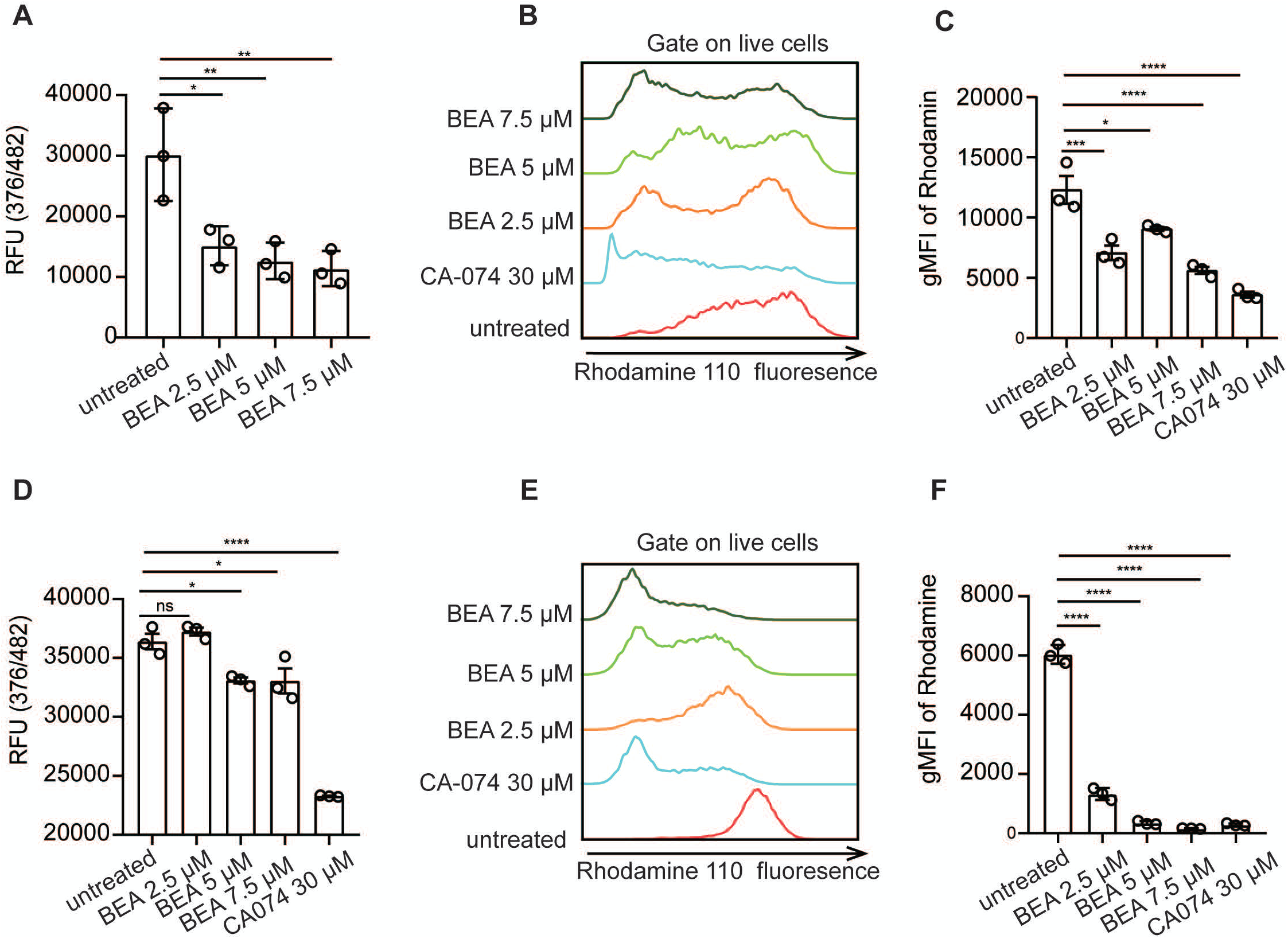
Inhibition of CTSB activity by BEA in mouse BMDCs. **(A)** 1x10^6^ BMDCs from C57BL/6N mice were stimulated with the indicated concentrations of BEA for 16 h, CTSB activity was measured in cell lysates. **(B, C)** 5x10^5^ BMDCs from C57BL/6N mice were stimulated with the indicated concentrations of BEA or the known CTSB inhibitor CA-074 for 16 h, CTSB activity was detected in BMDCs by flow cytometry. **(D)** 1x10^6^ iDCs were stimulated with the indicated concentrations of BEA for 16 h, CTSB activity was measured in cell lysates. **(E, F)** 5x10^5^ iDCs were stimulated with the indicated concentrations of BEA or CA-074 for 16 h, CTSB activity was detected in iDCs by flow cytometry. Data is shown as means ± SEM. Results shown are representative of two to three independent experiments. \**p* < 0.05, \*\**p* < 0.01, \*\*\*\**p* < 0.0001, ns: not significant.

### STD NMR spectroscopy of BEA binding to CTSB

To corroborate that the observed biological effects of BEA are mediated by direct interaction with CTSB, we performed saturation transfer difference (STD) NMR experiments using human CTSB. This technique allows identification of the chemical moieties of a ligand that interact directly with a target protein by measuring the shift in proton resonances that are in close proximity (41). The results show a strong increase in the signal of aromatic protons belonging to one benzene ring and the alkyl protons in one isopropyl group (**Figure 2**), indicating that this region of the molecule is directly binding to human CTSB.

**Fig 2.**
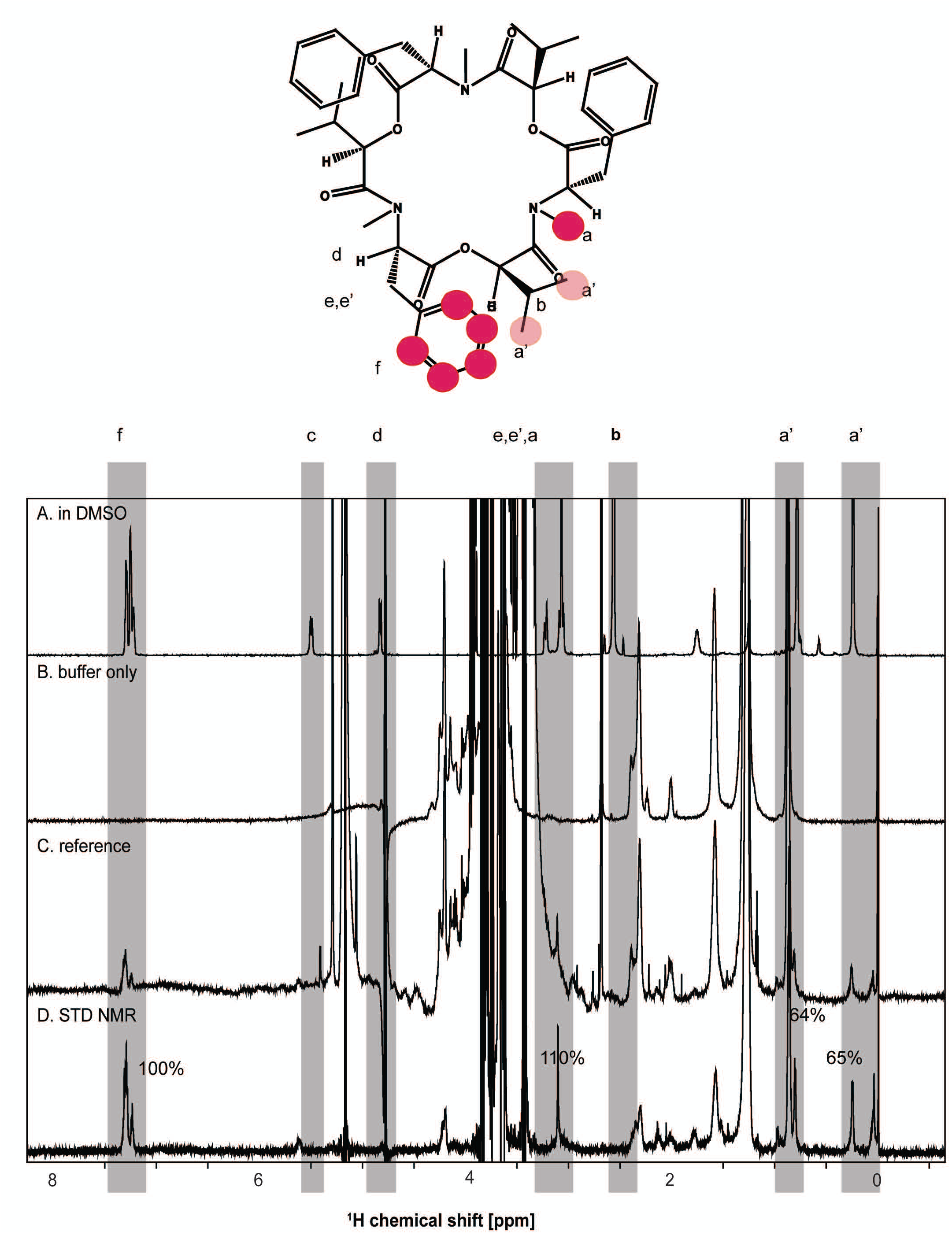
STD NMR spectra of BEA in the presence of hu CTSB. Chemical structure of BEA along with proton assignment and NMR spectra. **(A)** 1D ^1^H NMR of 250 µM BEA measured (256 scans) in 100 % (v/v) DMSO_d6_. **(B)** Control experiments of only buffer (128 scans), containing 25 mM oxalic acid/sodium oxalate, 5% (v/v) DMSO_d6_, pH 5.0, 10% (v/v) D_2_O, and 0.1% (v/v) Tween 20. **(C)** Reference 1D NMR spectrum (128 scans) and **(D)** STD NMR spectrum (4928 scans) of BEA in the presence of human CTSB, respectively. Protons making close contacts with the protein interaction site show significant enhancements in their resonances. The intensity of the STD effect of the aromatic protons was set to 100% as a reference, and relative intensities were determined. In particular, aromatic (f) and methyl (a, a’^-^’’) proton groups show strong signal enhancements. For all spectra, the scale for chemical shifts is shown at the bottom. The intensity of the spectra (scale of the y-axis) is adjusted for clarity.

### Enzyme kinetics of CTSB in the presence of BEA

Since the binding of BEA could occur either via the active site of CTSB or an allosteric site, we performed a kinetic characterization of the enzyme at different concentrations of the inhibitor, with the aim of determining whether BEA acts as a competitive, non-competitive, or uncompetitive inhibitor. Human CTSB shows a classical Michaelis-Menten behavior in the absence of an inhibitor (**Figure 3A**). As the inhibitor concentration increases, the maximal enzyme velocity *V*_max_ decreases while the K_M_ value remains similar within the experimental uncertainty (**Table 1**), suggesting that BEA does not act as a competitive inhibitor of the enzyme (55), and, therefore, does not bind directly to the active site of huCTSB.

**Fig 3.**
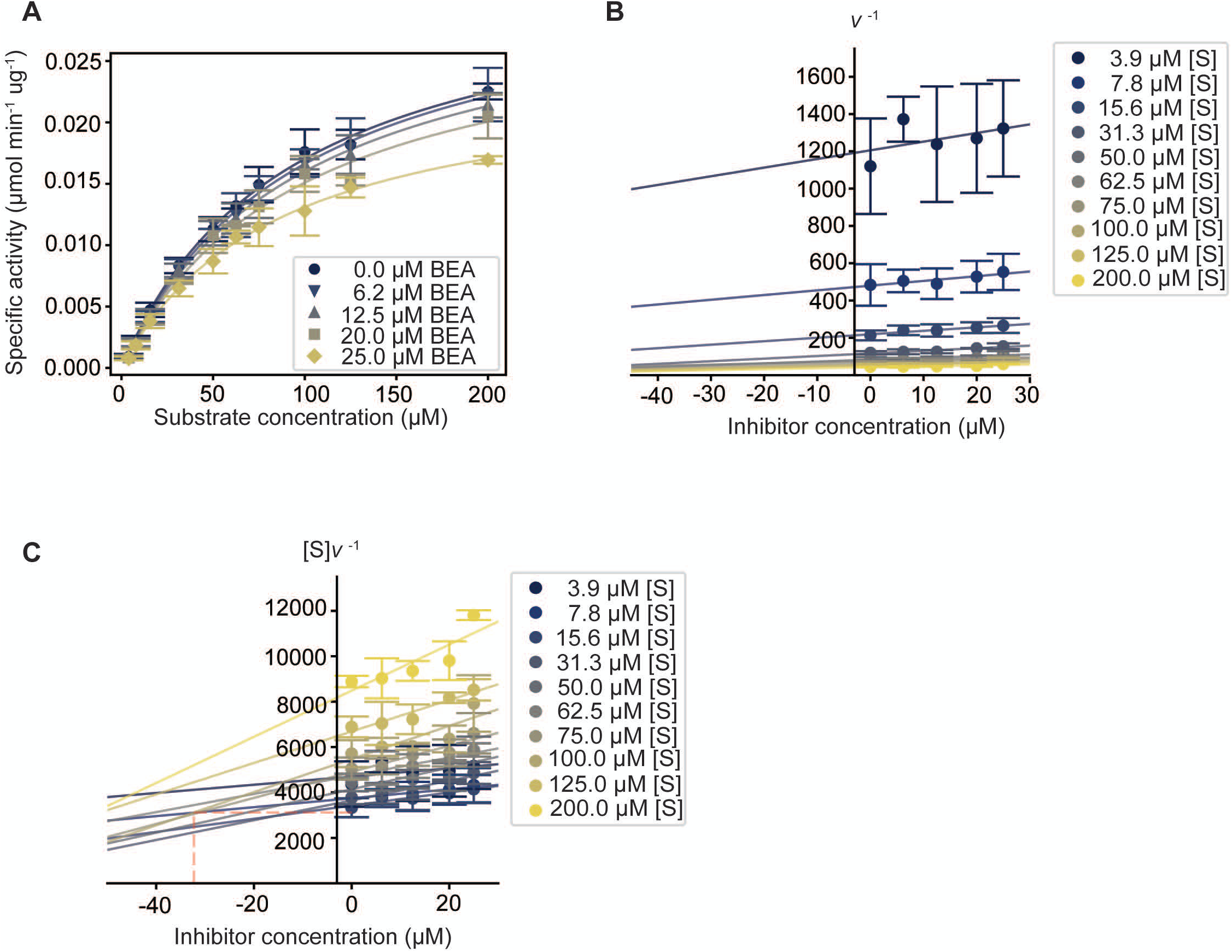
Kinetic characterization of BEA inhibition reveals an uncompetitive mechanism. **(A)** Michaelis-Menten curves for human CTSB in the presence of increasing concentrations of BEA. The lines show the model fit. **(B)** Dixon plot for the kinetic experiment shown in **(A)**. The lines show the linear fit. **(C)** Substrate over enzyme velocity plot for the kinetic experiment shown in **(A)**. The red dashed line shows the closest neighbor intersection point of the curves. The lines show the linear fit. In all plots, the error bars represent the SEM of a triplicate run.

**Table 1.** Kinetic parameters derived from Figure 3A.

| Inhibitor concentration ($\mu\text{M}$) | $K_m$ ($\mu\text{M}$) | $V_{\max}$ ($\text{pmol min}^{-1} \mu\text{g}^{-1}$) |
| --- | --- | --- |
| 0 | $93.9 \pm 6.5$ | $33.02 \pm 0.001$ |
| 6.25 | $99.1 \pm 8.1$ | $33.08 \pm 0.001$ |
| 12 | $99.5 \pm 7.0$ | $31.95 \pm 0.001$ |
| 20 | $98.3 \pm 13.3$ | $29.92 \pm 0.002$ |
| 25 | $86.2 \pm 5.2$ | $24.39 \pm 0.001$ |

To clarify whether BEA acts via an uncompetitive or mixed inhibition mechanism, we employed the graphical discrimination method proposed by Cornish-Bowden (42), which consists of plotting the reciprocal of the velocity (1/v) versus the concentration of the inhibitor ([I]), also known as Dixon plot (56), and the substrate concentration over the velocity ([S]/v) versus the inhibitor concentration, at multiple values of the substrate concentration. For a purely non-competitive inhibitor, both 1/v vs [I] and [S]/v vs. [I] will give straight lines converging in the negative region of the x-axis with y-values close to zero. For a mixed inhibitor, the lines will also converge in the negative region of the x-axis, but with y ≠ 0. Finally, an uncompetitive inhibitor will show parallel lines for the Dixon plot whereas the curves will intersect in the fourth quadrant of the [S]/v vs. [I] plot.

**Figure 3B and 3C** show the plots 1/v or [S]/v versus [I], respectively. In the first plot (**Figure 3B**), the curves do not intersect each other. In the second plot (**Figure 3C**), there are multiple intersections at x-values < 0 and y-values > 0, which strongly suggests that BEA behaves as an uncompetitive inhibitor. The x-value of the intersection point between the curves corresponds to the negative value of the inhibition constant (-*K*_i_). However, the experimental noise in the assays did not allow for the direct derivation of the K_i_ value. Therefore, we approximated the closest neighbor intersection point from all the curves by finding the value that minimizes the total distance to the multiple intersecting points (shown as the cross between the red dashed lines in **Figure 3C**). This results in an estimated *K*_i_ value of 44.3 µM. From equation (2), it can be deduced that the y-value of the intersection point of the curves is equal to K_M_/*V*_max_. As an internal check, we compared the experimental value of K_M_/*V*_max_ in the absence of inhibitor (2843.73 min µg) to the y-value of the closest neighbor intersection point (3116.68 min µg), revealing a difference of 9.15%, which indicates that our approximation yields a *K*_i_ of equal order of magnitude.

Overall, the kinetic characterization of the inhibition of human CTSB reveals that BEA acts as an uncompetitive inhibitor.

### Structural modelling of CTSB and BEA interactions

An uncompetitive inhibition mode implies that the inhibitor binds to an already established enzyme-substrate complex, ruling out the possibility of BEA binding directly to the active site of human CTSB. Apart from the active site, a regulatory site involved in the heparin-mediated activation has been described (57), but no inhibitory allosteric sites have been reported. To identify alternative putative binding sites for BEA, we carried out molecular dynamics simulations (MD) using multiple starting structures of human CTSB in the presence of two co-solvent probes that mimic the key chemical moieties of BEA found to be in direct contact with the enzyme by STD-NMR, benzene and isopropanol. Performing MD simulations in the presence of small molecular probes has been shown to be a powerful method to identify previously undescribed ligand binding sites (58, 59), as smaller probes tend to accumulate in the protein pockets that have the molecular features necessary for the binding of larger ligands. A large loop adjacent to the active site of CTSB, known as the occluding loop, displays a high degree of conformational variability (60). To avoid biasing our results by the starting conformations of the occluding loop, we performed the co-solvent MD simulations using four different starting conformations modeled from different crystal structures. **Figure 4** shows the regions where benzene (orange densities) and isopropanol (cyan densities) accumulate preferentially during the simulation for each one of the starting conformations of human CTSB. Besides small patches of interacting spots on the surface of the protein, for all the starting structures, the largest co-localization of probes occurs adjacent to the occluding loop near the active site.

**Fig 4.**
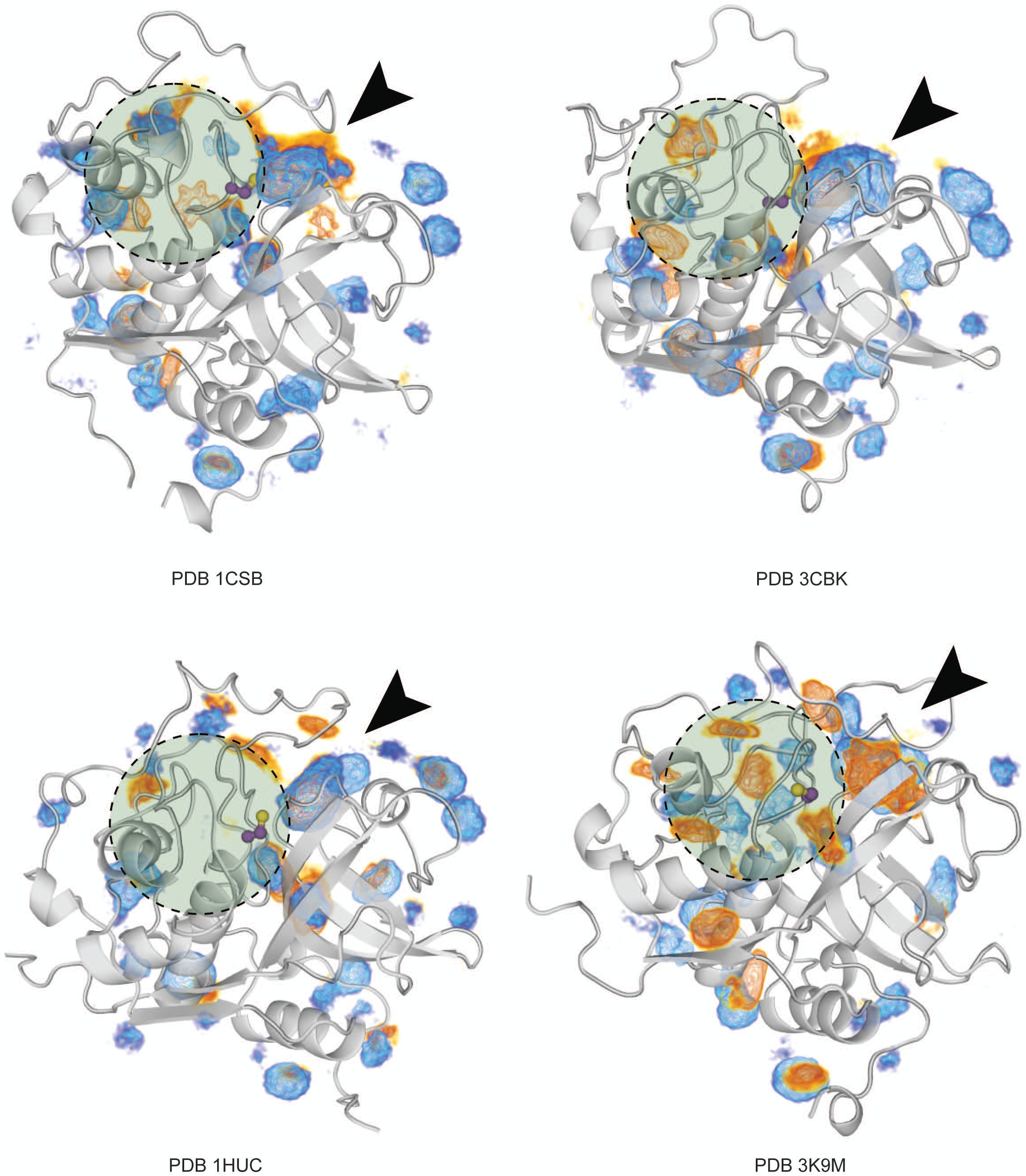
Co-solvent MD simulations reveal a common accumulation site adjacent to the catalytic site. Molecular dynamics simulations were started using four different initial conformations of human CTSB, each labelled according to the PDB structure that was used as template. The accumulation of co-solvent probes is shown as blue and orange densities for isopropanol and benzene, respectively. The volumes are shown in the same scale across panels. The largest accumulations are highlighted with arrow heads. The catalytic cysteine is shown as purple stick and highlighted with a green circle.

Next, we investigated if other known inhibitors of cathepsin could target the edge of the substrate binding site cleft adjacent to the occluding loop, as suggested by the MD simulations. Most of the small molecule inhibitors of human CTSB are covalent inhibitors that directly target the catalytic serine in the active site. Nonetheless, human CTSB as well as other proteases are expressed as autoinhibited zymogens that undergo activation under certain pH and redox conditions (61). The zymogen form includes a large N-terminal loop that covers the entirety of the substrate binding cleft and inhibits enzyme activity. Notably, the zymogen form of human CTSB (PDB ID 3PBH) contains an autoinhibitory peptide with a hot spot of Phe and Val residues interacting with the edge of the substrate binding cleft near the occluding loop, very similar to the interaction site we identified through MD simulations (**Figure 5A**). Previous work has reported that BEA can inhibit other papain-like enzymes such as papain and cathepsin V (62). Therefore, we assessed whether the autoinhibitory peptides of these enzymes also display a similar interaction motif in the same spot. **Figure 5A** shows that the autoinhibitory peptides of papain and cathepsin V have a Phe-Val-rich motif interacting directly with the edge of the substrate binding cleft, in the same position as observed for human CTSB.

**Fig 5.**
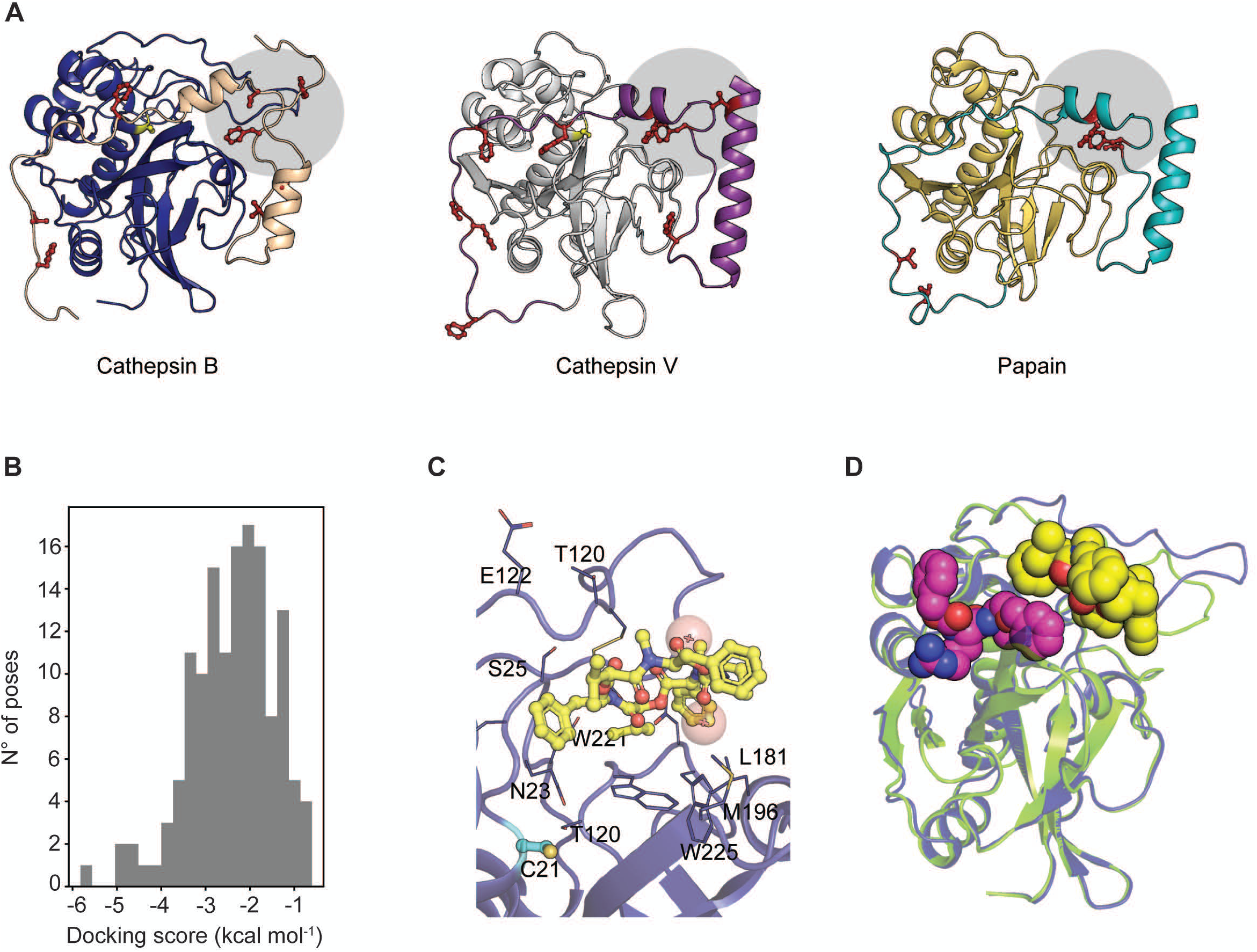
Identification of a putative BEA binding site using molecular docking. **(A)** Different proteins inhibited by BEA (Cathepsin B: blue, Cathepsin V: white, Papain: yellow) display Phe-Val-rich regions in their proenzyme autoinhibitory peptides (Cathepsin B: beige, Cathepsin V: purple, Papain: cyan). The residues are shown as red sticks, the region is highlighted with a grey circle. **(B)** Docking score distribution for the generated binding poses of BEA. **(C)** Binding mode consistent with the placement of the autoinhibitory peptide. BEA is shown as yellow sticks, nearby residues are shown as blue sticks, the catalytic cysteine is shown in cyan. The center of mass of the side chains of the autoinhibitory peptide are shown as red spheres. **(D)** Superimposition of the binding mode shown in **C** (protein shown as blue cartoon, BEA shown as yellow van der Waals spheres), against the structure of rat CTSB bound to the tripeptide-mimetic covalent inhibitor benzyloxycarbonyl-Arg-Ser(O-Bzl) chloromethylketone (PDB ID 1THE) (protein shown as green cartoon, inhibitor shown as purple van der Waals spheres).

Finally, we tested whether BEA could interact with human CTSB without overlapping directly with the catalytic site. For this, we performed molecular docking focusing the search of putative binding poses in the region where the Phe-Val hot spot of the autoinhibitory peptide is located. Using the autoinhibited conformation of human CTSB without the N-terminal inhibitory peptide as a target structure, we generated 141 binding poses using Glide SP (54). Overall, the docking scores of the generated poses are low, suggesting weak binding (**Figure 5B**). Within the top five poses with the highest score is one binding mode in which BEA is placed such that its benzene ring and isopropyl group occupy the same region as the Phe-Val hot spot of the autoinhibitory peptide and are far apart from the catalytic cysteine residue (**Figure 5C**). To verify if BEA could bind to this region without obstructing access to the catalytic center of the enzyme, we superimposed the predicted binding mode with the crystal structure of a covalent peptidomimetic inhibitor (**Figure 5D**). As evidenced by the van der Waals radius of the atoms of each ligand, the volume of both molecules does not overlap, suggesting that BEA could bind to human CTSB in a manner consistent with the uncompetitive inhibition mechanism.

## Discussion

For BEA, various biological functions have been reported including anti-cancer (24, 63–66), anti-inflammatory (27, 28), anti-viral (25, 26), and anti-microbial properties (67). Furthermore, an immune-activating function of BEA was identified in that BEA activates BMDCs via targeting TLR4, inducing inflammatory cytokine production (33). In silico prediction tools showed that CTSB is a potential target of BEA. Recently, Silva et al. indicated that BEA acts as a potent inhibitor against human CTSB in cell-free enzymatic activity assays. Here, we revealed that CTSB is active in BMDCs and that BEA significantly inhibits mouse CTSB activity at 2.5 μM in BMDCs after 16 h of stimulation. A similar inhibition of human CTSB activity by BEA was observed in human iDCs, and a significant inhibition was also observed at 2.5 μM. Moreover, through STD-NMR and enzyme kinetics characterizations, we established that BEA binds to CTSB and inhibits it, without displacing the substrate from the active site. Finally, computational structural analysis of CTSB revealed a putative binding site for BEA, where instead of acting as a substrate mimic, it mimics a motif found within the inhibitory propeptide region.

CTSB is the first and currently best-characterized member of the C1 family of papain-like, lysosomal cysteine peptidases (68). It is widely expressed in most cell and tissue types, and is mainly localized in endo/lysosomes and can be released into cytosol in inflammatory diseases (69). It was shown that elavated expressions of CTSB are observed in a variety of human diseases, such as cancer, neurodegeneration and immunological and developmental disorders. Increasing evidence of the pathophysiological roles and substrates of CTSB was revealed by the establishment of a CTSB knockout mouse (70). Gonzalez-Leal et al. showed that CTSB deficiency can lead to increased MHC-II molecule expression and can induce IL-12 production by LPS-treated BMDCs (71). This suggested that BEA might induce increased IL-12 production after inhibition of CTSB activity or deletion of CTSB in BMDCs. Further studies need to be done to verify whether BEA can induce more IL-12 production by BEA-treated BMDCs after inhibition of CTSB activity or deletion of CTSB.

We used STD-NMR to map the BEA epitope binding to human CTSB. The results showed that human CTSB interacts through the exposed side chains of phenylalanine and valine residues. Given that these residues are identical to amino acids of the peptide substrate known to mediate the interaction with the active site of human CTSB (75), one might speculate that BEA acts as a competitive inhibitor. However, the kinetic characterization clearly showed a decrease of *V*_max_, hence, discarding the possibility of BEA being a pure competitive inhibitor. Furthermore, graphical analysis via the Dixon plot and the [S]/v vs. [I] plot revealed that BEA acts as an uncompetitive inhibitor under the experimental conditions, which prompted the search for an alternative binding site. Although this inhibition mechanism is rare, it is sometimes explained by supposing that the inhibitor can bind the ES complex but not the free enzyme (76).

MD simulations in the presence of co-solvent probes did not reveal a novel marked allosteric site. Instead, the probes accumulated at the edge of the substrate binding cleft, which does not not contradict the proposed mechanism. Since CTSB has a large substrate binding cleft delimited by flexible loops that can accommodate a wide variety of substrates and inhibitors (60, 75), it is conceivable that human CTSB can bind a BEA molecule within the substrate binding cleft, while there is still enough space in the catalytic center to allow for the peptidase activity to take place. To further support the idea that the edge region of the binding cleft of human CTSB effectively interacts with BEA, we analyzed the contacts that the autoinhibitory peptide of the proenzymatic form of human CTSB forms with the rest of the structure. Since the autoinhibitory peptide contains pharmacophoric moieties required to abort human CTSB function, we hypothesized that there is a common binding subregion for both. Structural studies of the proenzymatic form of human CTSB showed that there is indeed a Phe-Val-rich region in the autoinhibitory peptide. Furthermore, by comparing different structures, we showed that the presence of this Phe-Val rich hot spot is common to the enzymes inhibited by BEA.

To test whether the identified site could accommodate BEA, we performed molecular docking focusing pose generation onto this region. Our results yielded a single pose, among the top-ranked ones, that is consistent with the space occupied by the autoinhibitory peptide. Furthermore, the observed STD-NMR signal shows that the interaction between BEA and CTSB occurs through the Phe and Val side chains of the peptide, which is consistent with the proposed binding mode. Considering that BEA acts as an uncompetitive inhibitor, conformational rearrangements induced by the presence of the substrate in the active site are required for the binding of BEA (75). Moreover, it has been shown that different inhibitors can trigger different conformational changes in CTSB due to its high flexibility (60). In our docking study, we used the autoinhibited conformation of human CTSB as target structure, as it might have the occluding loop in a position to allow the accommodation of BEA. However, molecular docking can be sensitive to structural variations of the protein target (77). Thus, it is likely that the overall low scores of the poses resulted from using a conformational state that is not similar enough to the conformation induced by the combined presence of the substrate and BEA. Further experimental structural studies are required to unravel the conformational dynamics of BEA binding to human CTSB, and how this hinders the enzyme function.

Taken together, our results show that BEA can suppress mouse and human CTSB activity in DCs. Moreover, we define the inhibition mode by showing that BEA can directly interact with human CTSB as an uncompetitive inhibitor. Molecular simulations and docking suggest a binding site at the edge of the active site, which is concordant with known structural data of other CTSB inhibitors. These first insights into the molecular mechanism of how BEA inhibits human CTSB should help further evaluating BEA as a candidate for cancer therapy.

## Author contributions

St.S. conceived and supervised the study together with H.G., and they reviewed and edited the manuscript. X.Y. designed and performed the experiments, analyzed the results, and wrote the manuscript. P.M. designed and performed enzymatic assays, molecular simulations, and docking analyses and wrote the manuscript. M.H. performed NMR experiments and wrote the manuscript. A.V. performed CTSB assays and So.S. generated BMDCs. S.O. and S.M. performed the measurements of CTSB assays. A.H. and J.Q. wrote the manuscript. D.F. and M.U. gave suggestions on the project.

## Acknowledgement

This work was funded by the Deutsche Forschungsgemeinschaft (German Research Foundation) DFG-270650915/GRK2158. We are grateful for computational support and infrastructure provided by the “Zentrum für Informations- und Medientechnologie” (ZIM) at the Heinrich Heine University Düsseldorf and the computing time provided by the John von Neumann Institute for Computing (NIC) to H.G. on the supercomputer JUWELS at Jülich Supercomputing Centre (JSC) (user ID: VSK33).

